# NSD2 promotes cell durotaxis and drives the transition from PKD to tubulocystic renal cell carcinoma through integrin/FAK/AKT signaling

**DOI:** 10.1101/2024.10.21.619559

**Authors:** Wenxin Feng, Ningyuan Liu, Changwei Liu, Hanyu Rao, Zhuo Chen, Wei Zhang, Yue Xu, Rebiguli Aji, Ziyi Wang, Wei-Qiang Gao, Li Li

**Affiliations:** 1 State Key Laboratory of Systems Medicine for Cancer, Renji-Med X Clinical Stem Cell Research Center, Ren Ji Hospital, School of Medicine and School of Biomedical Engineering, Shanghai Jiao Tong University, Shanghai, 200127, China; 2 School of Biomedical Engineering and Med-X Research Institute, Shanghai Jiao Tong University, Shanghai, China

**Keywords:** NSD2, Integrin, FAK signaling, TCRCC, H3K36me2

## Abstract

Renal cell carcinoma (RCC) is one of the most common malignancies in the urinary system. NSD2 is an H3K36-specific di-methyltransferase which has been reported to participate in diverse biological processes and human tumors. However, its role in RCC remains unclear. Here, we found that NSD2 is highly expressed in RCC, which is associated with poor survival in RCC patients. NSD2 facilitates the transition from *Myc*-induced polycystic kidney disease to tubulocystic renal cell carcinoma (TCRCC), which is a rare RCC subtype with distinctive clinicopathologic and genetic characterizations. The mice with kidney-specific overexpression of MYC and NSD2 (KMN) display severe cyst burden at only 6 weeks of age, and develop into TCRCC at 12 weeks of age. Mechanistically, NSD2 transcriptionally upregulates the expressions of integrins (*Itga4* and *Itga11*), to further activate the FAK/AKT pathway. In addition, we found that NSD2 enhances the cell proliferation on the stiff matrix of PEGDA hydrogel. Moreover, inhibition of FAK signaling relieves the symptoms of KMN mice, and significantly rescues the enhanced cell proliferation caused by NSD2 overexpression in vitro. Together, our findings highlight an epigenetic mechanism by which NSD2 regulates TCRCC tumorigenesis through the integrin/FAK/AKT pathway. This study may also pave the way for the development of targeted, patient-tailored therapies for TCRCC patients with *NSD2* amplification or high expression.

## Introduction

Tubulocystic renal cell carcinoma (TCRCC) is a rare renal cell carcinoma (RCC) subtype which is classified as a distinct kind of tumor of the kidney in the WHO until 2016 (1). TCRCC is characterized by aggressive growth of the cysts, showing variably sized cystically dilated tubules (2,3). It has been reported that only less than 100 cases of pure TCRCC have been identified, thus the mechanism of TCRCC tumorigenesis remains to be explored (4).

Increasingly studies have demonstrated that RCC could be mediated by epigenetic modifiers (5). Exome sequencing studies also revealed frequent mutations in genes that encode histone-modifying and chromatin-modifying proteins, implicating epigenetic reprogramming as a key feature of renal carcinogenesis (6). For instance, overexpression of KDM5C leads to a poor prognosis of RCC patients, and loss of SETD2 accelerates RCC formation in *Myc*-driven polycystic kidney disease (PKD) mice (7,8). However, little is known about the relationship between TCRCC and epigenetic regulation.

NSD2 is a histone methyltransferase that specifically catalyzes the di-methylation of histone H3 at lysine 36 (H3K36me2) (9). The *NSD2* chromosomal translocation in multiple myeloma is associated with the overexpression of NSD2 and leads to a poor prognosis (10). Similar mutations also have been discovered in renal cell carcinoma and clinical samples show elevated H3K36me2 and altered gene activation (11,12). Previous studies have identified that NSD2 plays a significant role in epigenetic regulation of tumors, functioning in cell proliferation and metastasis (13). However, the role of NSD2 in RCC remains unclear.

As a receptor of integrin, focal adhesion kinase (FAK) can be phosphorylated in response to the uplevel of the integrin family, and leads to the activation of the downstream PI3K/AKT signaling pathways and mTOR phosphorylation (14). FAK is a highly tyrosine-phosphorylated protein, functioning as a major driver of tumorigenesis in various cancers (15). In addition, FAK can activate intracellular proteases such as calpain and extracellular matrix (ECM) metalloproteinases, involved in ECM-receptor interaction and cell proliferation (14,16,17). Moreover, evidence shows that downregulating the phosphorylation of FAK attenuates RCC tumorigenesis, highlighting the potential function of FAK on RCC (18).

In this study, we investigate the role of NSD2 in TCRCC tumorigenesis using transgenic mice with kidney-specific overexpression of MYC and NSD2. Our results demonstrated that overexpression of NSD2 leads to up-regulation of *Itga4* and *Itga11*, thus activating the integrin/FAK/AKT pathway. We found that overexpression of NSD2 can enhance cell proliferation on the stiff matrix. Therefore, these findings indicate that NSD2 plays a significant role in facilitating TCRCC tumorigenesis.

## Materials and Methods

### Mice

The *Ksp*^Cre^ mice (B6.Cg-Tg (Cdh16-cre)91Igr/J) were purchased from The Jackson Laboratory. NSD2-overexpressing mice (*Nsd2*^OE/+^) were generated as previously reported (19) and gifted by Prof. Jun Qin from Chinese Academy of Sciences. *Nsd2*^OE/+^ mice were mated with *Ksp*^Cre^ mice to generate *Ksp*^Cre^; *Nsd2*^OE/+^ mice. *Myc*^OE/+^ mice (20) were mated with *Ksp*^Cre^ mice to generate *Ksp*^Cre^; *Myc*^OE/+^ (KM) mice. *Nsd2*^OE/+^ mice were mated with KM mice to generate *Ksp*^Cre^; *Myc*^OE/+^; *Nsd2*^OE/+^ (KMN) mice. All the mice were born and maintained in the specific pathogen-free animal facility. All animals were randomized and exposed to the same environment. The mice that lose weight by more than 20% within 1 week will be euthanized and counted as death. All experimental procedures were approved by the Animal Ethics Committee of School of Biomedical Engineering & Med-X Research Institute, Shanghai Jiao Tong University.

### RNA isolation and quantitative RT-qPCR

Total RNA was extracted from cultured cells or tissues using an RNA extraction kit (BioTeke) according to the manufacturer’s protocol. The RNA was reverse transcribed using an RT kit (Takara), and the resulting cDNA was used for TB Green real-time PCR analysis. β-actin was used for normalization, and the data were presented as mean ± S.D. Student’s t-test was used to calculate the P value.

### Western blotting analysis and antibodies

Tissues and cells were harvested and lysed with RIPA buffer supplemented with protease and phosphatase inhibitors (MCE). The protein concentration was measured with the BCA Protein Assay (BioRad). The protein was separated by 8%–12% SDS-PAGE gels and transferred onto polyvinylidene fluoride membranes (Millipore). Membranes were blocked in 5% BSA in TBS for 1 hour at room temperature and then incubated with primary antibodies overnight at 4 ℃, washed in TBS containing 0.1% Tween20, incubated with horseradish peroxidase (HRP)-conjugated secondary antibody for 1 hour at room temperature, and developed by ECL reagent (Thermo Fisher Scientific). The primary antibodies used in this study were as follows: NSD2(Abcam, ab75359), H3K36me2 (Abcam, ab9049), H3 (Cell Signaling Technology, 9715), β-Tubulin (Cell Signaling Technology, 2146), E-cadherin (Cell Signaling Technology, 3195), Vimentin (Cell Signaling Technology, 5741), FAK (Cell Signaling Technology, 3285), p-FAK (Cell Signaling Technology, 3283), AKT (Cell Signaling Technology, 2920), p-AKT (Cell Signaling Technology, 4060).

### Histology and IHC staining

After washing kidney tissues with cold PBS, they were fixed in 4% paraformaldehyde at 4 ℃ overnight. Then, the tissues were dehydrated and embedded in paraffin. Sections (5 μm) were stained with hematoxylin and eosin (H&E). For IHC staining, sections were dewaxed and hydrated, followed by antigen retrieval in citrate buffer at high heat for 3 min and low heat for 12 min, cooled to room temperature, and washed three times with PBS. Endogenous peroxidase was quenched with 3% H2O2. Next, samples were blocked with 5% BSA at room temperature for 1 h. Then, samples were incubated with primary antibodies at 4 ℃ for 12-16 h. The primary antibodies used were NSD2 (Abcam, ab75359), H3K36me2 (Abcam, ab9049), AMACR (Abcam, ab140798), Vimentin (Cell Signaling Technology, 5741), Ki67 (Abcam, ab15580), CK19 (Abcam, ab15463), Collagen I (Cell Signaling Technology, 72026), ITGA11 (Abcam, ab198826), ITGA4 (Cell Signaling Technology, 8440), p-FAK (Cell Signaling Technology, 3283), p-AKT (Cell Signaling Technology, 4060), Collagen V (Abcam, ab7046). After washing three times with PBS, sections were incubated with HRP-conjugated secondary antibodies at room temperature for 1 h, followed by counterstaining with hematoxylin. Images were captured using a Leica microscope and staining intensity was calculated using ImageJ software.

### Poly (ethylene glycol) diacrylate (PEGDA) gel fabrication and characterization

Wipe the mirror with ethanol and then treat the mirror with Gel Slick solution. Mix PEGDA (MW∼575, Macklin) and 1 mg/ml Irgacure 2959 (2-hydroxy-4’-(2-hydroxyethoxy)-2-methylpropiophenone, Macklin) proportionally with PBS to produce a hydrogel precursor solution ranging from 4% to 10% (21–23). Drop 200 μl precursor solution on the mirror, and cover with 20 mm glass functionalized with 3-(Trimethoxysilyl) propyl methacrylate (Macklin), and polymerized with UV light (10 min, 365 nm, 60 J) to form 150-μm-thick hydrogel films. Remove the gel from the mirror and place it facing up in the appropriate size of the 12-well culture plate. Suck the MES solution on the surface of the water purification gel by the vacuum pump, and drop 400 μl 0.25 mg/ml sulfo-SANPAH with MES buffer to completely cover the gel surface, exposed to UV light (15 min, 365 nm, 60 J), and repeat twice. After rinsing with MES buffer for 5 min and then rinsing with HEPES buffer, 25 μg/ml human placenta type IV (Collagen IV, Corning) was prepared with HEPES buffer and incubated on the gel for 2.5 h, then incubated in 0.5 Methanolamine (Macklin) for 60 min to inhibit the function of unreacted sulfo-SANPAH. Rinse the gel with PBS buffer 4 times for 30 minutes. The PEGDA gel can be stored overnight at 4 ℃, rinsed twice with PBS buffer and sterilized under UV light for 2 hours before use.

### RNA sequencing and analyses

Kidney samples were obtained from 12-week-old KM and KMN mice. Total RNA was extracted from the samples by Trizol reagent (Invitrogen) separately. The RNA quality was checked by Agilent 2200 and kept at −80 ℃. The RNA with RIN (RNA integrity number) > 7.0 is acceptable for cDNA library construction. The cDNA libraries were constructed for each RNA sample using the TruSeq Stranded mRNA Library Prep Kit (Illumina, Inc.) according to the manufacturer’s instructions. Before read mapping, clean reads were obtained from the raw reads by removing the adaptor sequences and low-quality reads. The clean reads were then aligned to mouse genome (mm10) using the Hisat2. HTseq was used to get gene counts and RPKM method was used to determine the gene expression. We applied DESeq2 algorithm to filter the differentially expressed genes, after the significant analysis, P-value and FDR analysis were subjected to the following criteria: i) Fold Change>2 or < 0.5; ii) P-value<0.05. Gene ontology (GO) analysis was performed to facilitate elucidating the biological implications of the differentially expressed genes in the experiment. We downloaded the GO annotations from NCBI (http://www.ncbi.nlm.nih.gov/), UniProt (http://www.uniprot.org/) and the Gene Ontology (http://www.geneontology.org/). Fisher’s exact test was applied to identify the significant GO categories (P-value < 0.05). Pathway analysis was used to find out the significant pathway of the differentially expressed genes according to KEGG database. We turned to the Fisher’s exact test to select the significant pathway, and the threshold of significance was defined by P-value< 0.05.

### CUT&Tag assay

We separated primary renal tubular epithelial cells (PTECs) from 10-week-old *Nsd2*^OE/+^ and *Ksp*^Cre^; *Nsd2*^OE/+^ mice as previously described(24). The CUT&Tag assay was performed using anti-H3K36me2 (Abcam, ab9049) antibodies. The CUT&Tag assay was performed following the manual of the hyperactive pG-Tn5/pA-Tn5 transposase for CUT&Tag kit (Vazyme, TD903). DNA libraries were prepared according to the manufacturer’s instructions for the Trueprep index kit v2 (Vazyme, TD202). The DNA library was sequenced to an average of 20 million reads per sample. Raw reads were filtered to obtain high-quality clean reads by removing sequencing adapters, short reads (length <36 bp) and low-quality reads using Trimmomatic v0.38[2] (non-default parameters: SLIDINGWINDOW:4:15 LEADING:10 TRAILING:10 MINLEN:36). Then FastQC (with default parameters) is used to ensure high reads quality. The clean reads were mapped to the mouse genome (assembly GRCm38) using the Bowtie2 v2.3.4.1 (non-default parameters: -I 10 -X 700 --no-discordant --no-mixed --local --very-sensitive-local) software. Duplicate reads were then removed using picard MarkDuplicates. Peak detection was performed using the MACS v2.1.2 (non-default parameters: -f BAMPE -g hs/mm -q 0.05) peak finding algorithm with 0.05 set as the q-value cutoff. Annotation of peak sites to gene features was performed using the ChIPseeker R package.

### Statistical analysis

All experiments were performed using 3–15 mice or at least three independent repeated experiments. Unless otherwise indicated, data presented as the mean ± S.E.M. All statistical analyses were performed with GraphPad 8.0 software. Student’s t-test assuming equal variance was used, and two-way analysis of variance for independent variance. *p <0.05, **p<0.01, ***p<0.001 and ****p<0.0001. No data were excluded from the analyses.

## Results

### NSD2 expression increases in RCC patients and is associated with poor RCC prognosis

To explore a possible role of NSD2 in RCC, we analyzed *NSD2* mRNA expression in public datasets of RCC samples (using datasets from NCBI’s Gene Expression Omnibus: GSE53757). The results indicated that the level of *NSD2* mRNA was increased in RCC specimens compared with normal kidney controls (Fig. 1A). Moreover, the expression of *NSD2* was significantly increased in patients with advanced RCC (Fig.1B) (using datasets from NCBI’s Gene Expression Omnibus: GSE73731). We also performed immunohistochemistry (IHC) analyses using a pre-valuated NSD2 antibody to characterize the NSD2 protein expression in RCC samples and normal kidney samples (Fig. 1C). The quantification of IHC staining showed that the protein level of NSD2 in the RCC patients was significantly higher than that of normal kidney tissue (Fig. 1D).

**Fig. 1.**
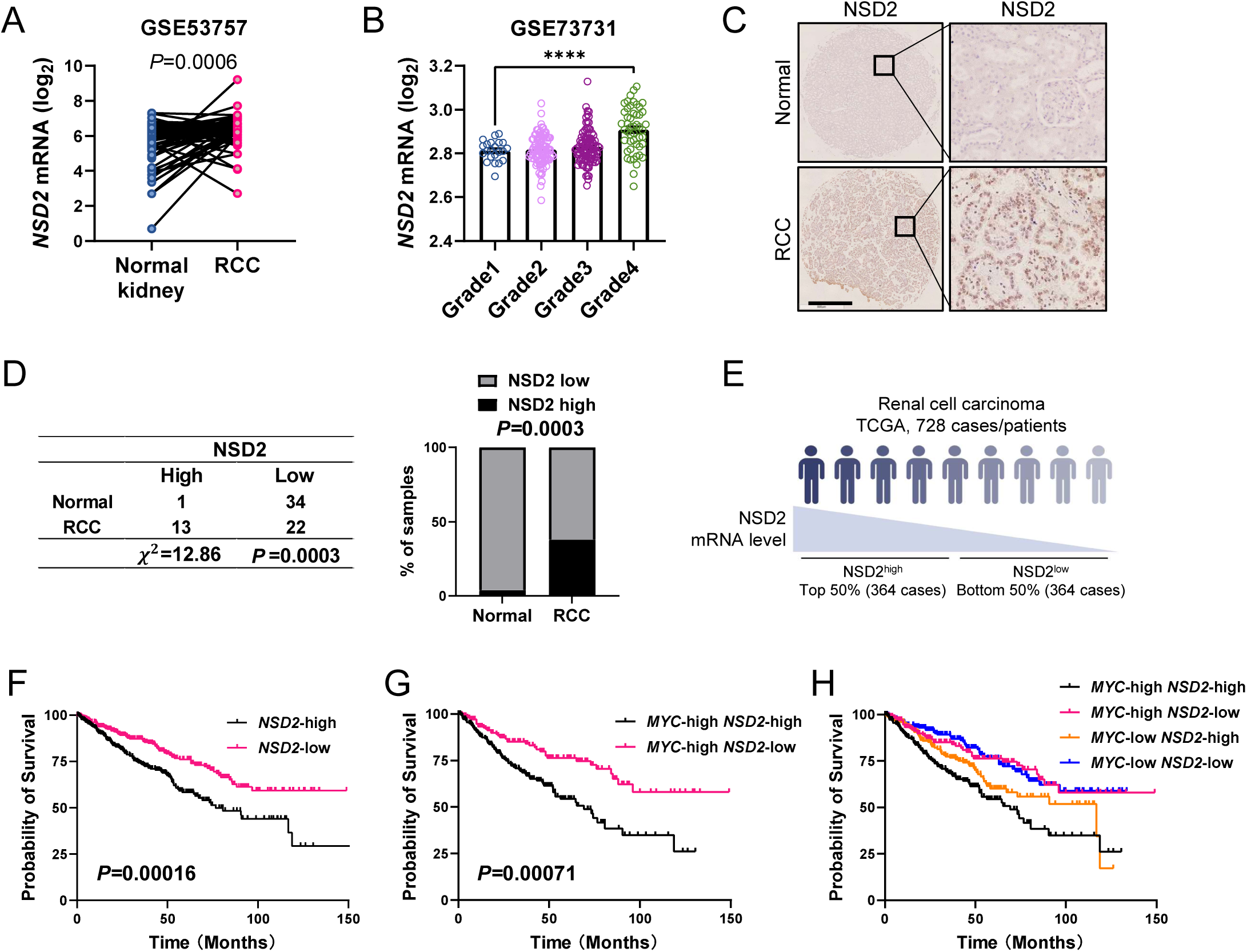
NSD2 expression increases in RCC patients and is associated with poor RCC prognosis. **(A)** The mRNA level of *NSD2* in normal kidney and RCC tissues from GEO database. **(B)** The mRNA level of *NSD2* in different grades of RCC patients from GEO database. **(C)** IHC analysis of NSD2 expression in normal (n = 35) and RCC (n = 35) tissues. Scale bars: 50 μm. **(D)** Statistics of high and low expression of NSD2 in normal (n = 35) and RCC (n = 35) tissues. **(E-F)** Survival curve of high (n = 364) and low (n = 364) *NSD2* expression from RCC patients in TCGA database. **(G-H)** Survival curve of high (n = 189) and low (n = 190) *NSD2* expression under the circumstances *MYC* high and low from RCC patients in TCGA database.

In addition, a Kaplan-Meier plot revealed a significant association between *NSD2* and overall survival. High expression of *NSD2* was correlated with significantly worse overall survival in RCC patients (Fig. 1E-F). Moreover, patients with both *MYC* and *NSD2* high expression have a worse clinical survival course (Fig. 1G-H). The collective results demonstrated that NSD2 expression increases in RCC patients and is associated with poor RCC prognosis.

### NSD2 promotes the transition from *Myc*-induced PKD to TCRCC

To investigate whether NSD2 overexpression could promote RCC tumorigenesis, we crossed *Nsd2*^OE/+^ mice with Ksp-Cre mice to obtain *Ksp*^Cre^; *Nsd2*^OE/+^ mice (Fig. S1A). Firstly, we found that the levels of NSD2 and H3K36me2 were significantly increased in *Ksp*^Cre^; *Nsd2*^OE/+^ mice (Fig. S1B-D). Histological examination of *Nsd2*^OE/+^ and *Ksp*^Cre^; *Nsd2*^OE/+^ mice at 12 months of age revealed no obvious abnormalities in the kidney (Fig. S1E-F). These results demonstrate that NSD2 overexpression is not sufficient to induce kidney tumors.

We next aimed to determine whether NSD2 is involved in the transition from *Myc*-induced PKD to RCC. We generated kidney-specific NSD2 overexpression mice together with oncogenic *Myc* mutation. *Myc*^OE/+^ mice were mated with *Ksp*^Cre^ mice to generate *Ksp*^Cre^; *Myc*^OE/+^ (KM) mice in C57BL/6 background. *Nsd2*^OE/+^ mice were mated with KM mice to generate *Ksp*^Cre^; *Myc*^OE/+^; *Nsd*2^OE/+^ (KMN) mice (Fig. S2A). The overexpression of NSD2 in the KMN mice was verified by using IHC, Western blotting, and Real-Time quantitative PCR (RT-qPCR) (Fig. 2A-B, S2B-C). In sharp contrast to age-matched KM mice, KMN mice did not typically survive past 20 weeks (Fig. 2C). KMN mice displayed a significant cystic phenotype at 6 weeks and were severely affected at 12 weeks, with significantly increased kidney volume and cystic index, as well as cysts number (Fig. 2D-G, S2D). The kidney of KMN mice is composed of numerous well-formed tubules and cysts which are lined by a layer of flat to hobnail cells with prominent nucleoli and separated by thin fibrous septa (Fig. 2E), which are typical features of TCRCC. Notably, compared with KM mice, KMN mice at 12 weeks of age showed significantly elevated CK19 and AMACR (Fig. 2H), which are strongly positive in all tubulocystic carcinoma cases (25–27). Besides, KMN mice displayed enhanced staining of Vimentin and Ki67 (Fig. 2H). Moreover, the levels of blood urea nitrogen (BUN) and creatinine were significantly increased in the blood of KMN mice (Fig. 2I).

**Fig. 2.**
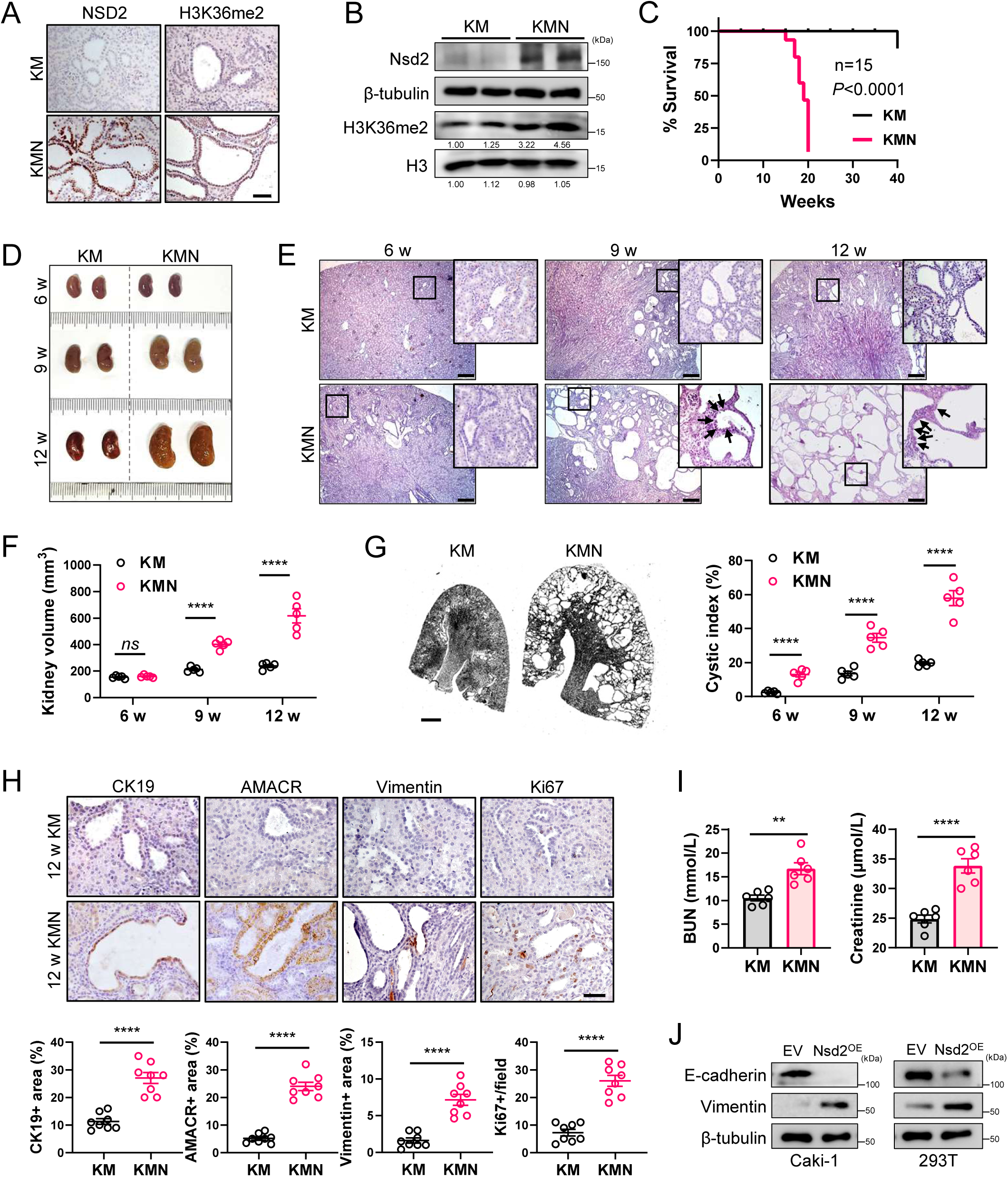
NSD2 promotes the transition from *Myc*-induced PKD to TCRCC. **(A)** IHC analysis of NSD2 and H3K36me2 levels in kidney tissues of KM and KMN mice (n = 5 per group). Scale bars: 50 μm. **(B)** Western blotting analysis of NSD2 and H3K36me2 levels in kidney tissues of KM and KMN mice. Experiments were repeated at least three times, with similar results. **(C)** Survival rate of KM and KMN mice (n = 15 per group). **(D)** Kidney images of KM and KMN mice at 6, 9, and 12 weeks. **(E)** Representative hematoxylin and eosin images of kidney tissue from 6-, 9-, and 12-week KM and KMN mice. The black arrows denote the hobnail cells. Scale bars: 50 μm. **(F)** Quantification of kidney volume (mm^3^) from indicated mice at 6, 9, and 12 weeks (n = 5 per group). **(G)** Histologic examination of kidneys from KM and KMN mice. Scale bars: 2 mm. Quantification of cystic index (%) in kidney tissue from indicated mice at 6, 9, and 12 weeks is on the right (n = 5 per group). **(H)** IHC staining and quantification of CK19, AMACR, Vimentin positive area, and Ki67 positive cells per field from indicated mice at 12 weeks (n = 8 per group). Scale bars: 50 μm. **(I)** Blood urea nitrogen and creatinine levels from indicated mice at 12 weeks (n = 6 per group). **(J)** Western blotting analysis of E-cadherin and Vimentin expressions in EV and Nsd2-OE cells (Caki-1 and 293T derived). Experiments were repeated at least three times, with similar results.

To further explore the effect of NSD2 overexpression, empty vector (referred to as EV) and NSD2 overexpression (referred to as Nsd2-OE) plasmids were transfected respectively in Caki-1 cells. NSD2 overexpression promoted colony formation ability in vitro (Fig. S2E). What’s more, Nsd2-OE cells showed drastically impaired expression of epithelial marker E-cadherin and elevated expression of mesenchymal marker Vimentin (Fig. 2J), indicating that NSD2 overexpression enhanced epithelial-mesenchymal transition.

Together, our results demonstrated that NSD2 promotes the transition from *Myc*-induced PKD to TCRCC.

### NSD2 activates the ECM-receptor interaction and FAK/AKT pathway

To gain mechanistic insight into how NSD2 promotes renal tumorigenesis, we performed RNA sequencing (RNA-seq) with renal tissues from KM and KMN mice (Fig. 3A-B). The standard for screening differentially expressed genes was FC>2 and P-value<0.05 was a significant differential gene. We identified 1657 differentially expressed genes, including 592 upregulated genes and 1065 downregulated genes. KEGG pathway analysis showed that NSD2 overexpression significantly enriched the genes associated with ECM-receptor interaction, focal adhesion, and PI3K-AKT signaling pathway (Fig. 3C). To confirm that overexpression of NSD2 activates these signaling pathways, RT-qPCR was performed and the results showed that the expression levels of genes in these related pathways were significantly changed in KMN mice (Fig. 3D).

**Fig. 3.**
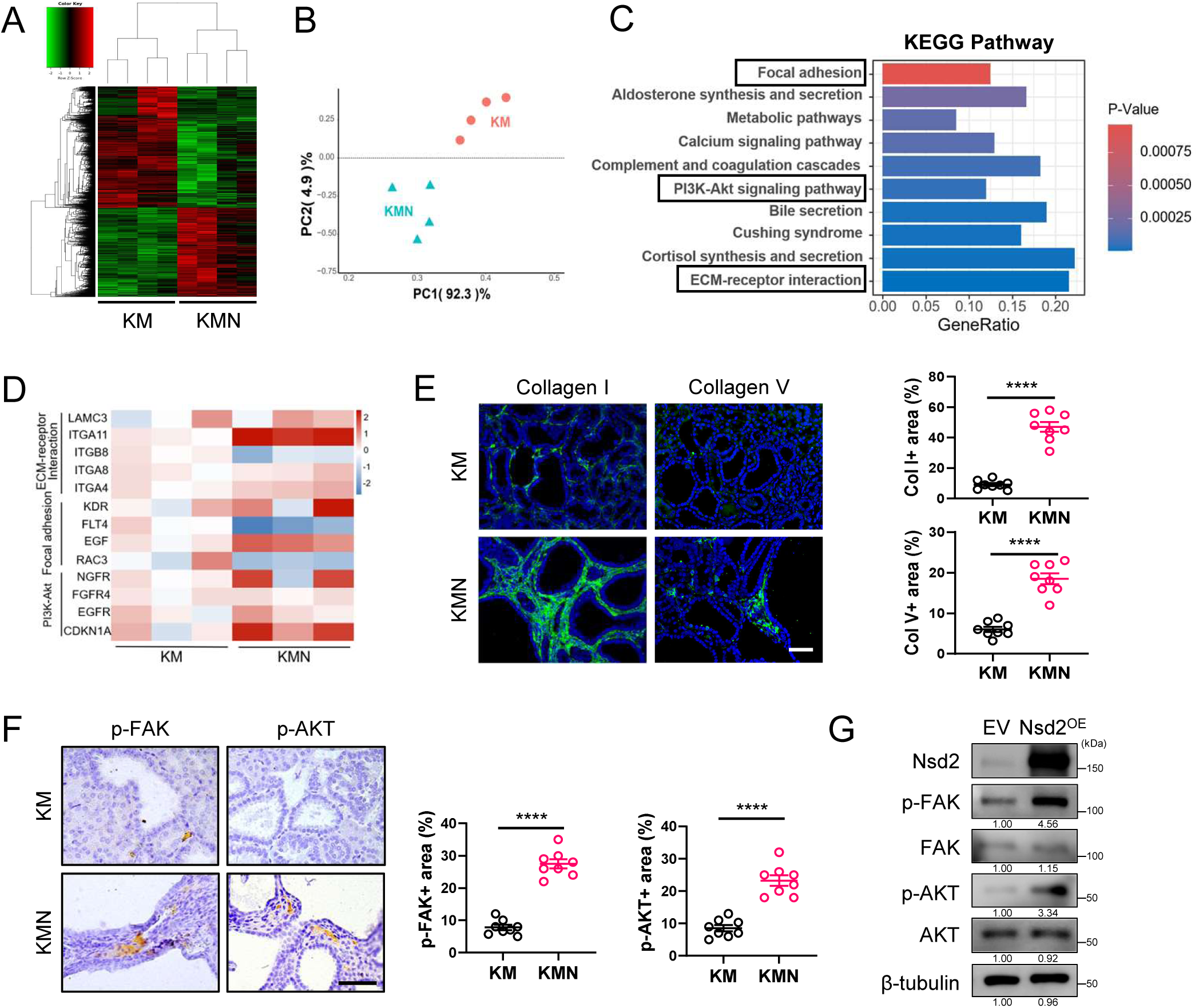
NSD2 activates the ECM-receptor interaction and FAK/AKT pathway. **(A)** Heat map of RNA-seq data to compare the gene expression of kidney tissue derived from KM and KMN mice (n = 4 per group). **(B)** Principal component analysis (PCA) scores plot indicating discrimination between KM and KMN mice (n = 4 per group). **(C)** KEGG pathway analysis of gene expression changes in RNA-seq data (KM and KMN mice). **(D)** RT-qPCR analysis of ECM-receptor interaction, Focal adhesion, and PI3K/AKT signaling related genes mRNA levels in kidney tissues of KM and KMN mice (n = 3 per group). **(E)** IF analysis of Collagen I and Collagen V expressions in KM and KMN mice at 12 weeks (n = 8 per group). Quantification of Collagen I and Collagen V positive area is shown on the right. Experiments were repeated at least three times, with similar results, and representative images are shown. Scale bars: 50 μm. **(F)** IHC staining and quantification of p-FAK and p-AKT expressions in KM and KMN mice at 12 weeks (n = 8 per group). Experiments were repeated at least three times, with similar results, and representative images are shown. Scale bars: 50 μm. **(G)** Western blotting analysis of NSD2, FAK, p-FAK, AKT, and p-AKT expressions in EV and Nsd2-OE cells (Caki-1 derived). Experiments were repeated at least three times, with similar results.

It has been reported that increased ECM stiffness leads to integrin clustering and cytoskeletal reinforcement. Phosphorylation of FAK plays a pivotal role in ECM stiffness-induced cell tension and AKT activation (28).

To further confirm that NSD2 activates the ECM-receptor interaction pathway, immunofluorescence (IF) staining was performed and the results showed that more Collagen I and Collagen V were expressed in KMN mice (Fig. 3E). In addition, the phosphorylation levels of FAK and AKT were significantly upregulated in KMN mice, indicating that FAK/AKT signaling was activated (Fig. 3F). The Western blotting also confirmed the activation of the FAK/AKT signaling pathway in Nsd2-OE cells (Fig. 3G).

Together, these results proved that NSD2 overexpression activates the ECM-receptor interaction and FAK/AKT pathway.

### NSD2-mediated H3K36me2 facilitates the expression of integrins

Considering that NSD2 is an H3K36-specific di-methyltransferase, CUT&Tag analysis of H3K36me2 was performed to further understand the underlying mechanisms. In line with our hypothesis, H3K36me2 peaks were enriched around the transcription area (Fig. 4A, S3). Focusing on genes with increased H3K36me2, KEGG pathway analysis was performed and the results showed that these genes were associated with ECM-receptor interaction and focal adhesion (Fig. 4B). To correlate the chromatin binding with the transcriptional regulation, we integrated the CUT&Tag data with the expression profile. The Venn diagram indicated that 231 genes showed direct H3K36me2 occupancies and expression changes upon the NSD2 overexpression (Fig. 4C). KEGG pathway analysis showed that the overlapping genes were still closely related to the ECM-receptor interaction, focal adhesion, and PI3K-AKT signaling pathway (Fig. 4D).

**Fig. 4.**
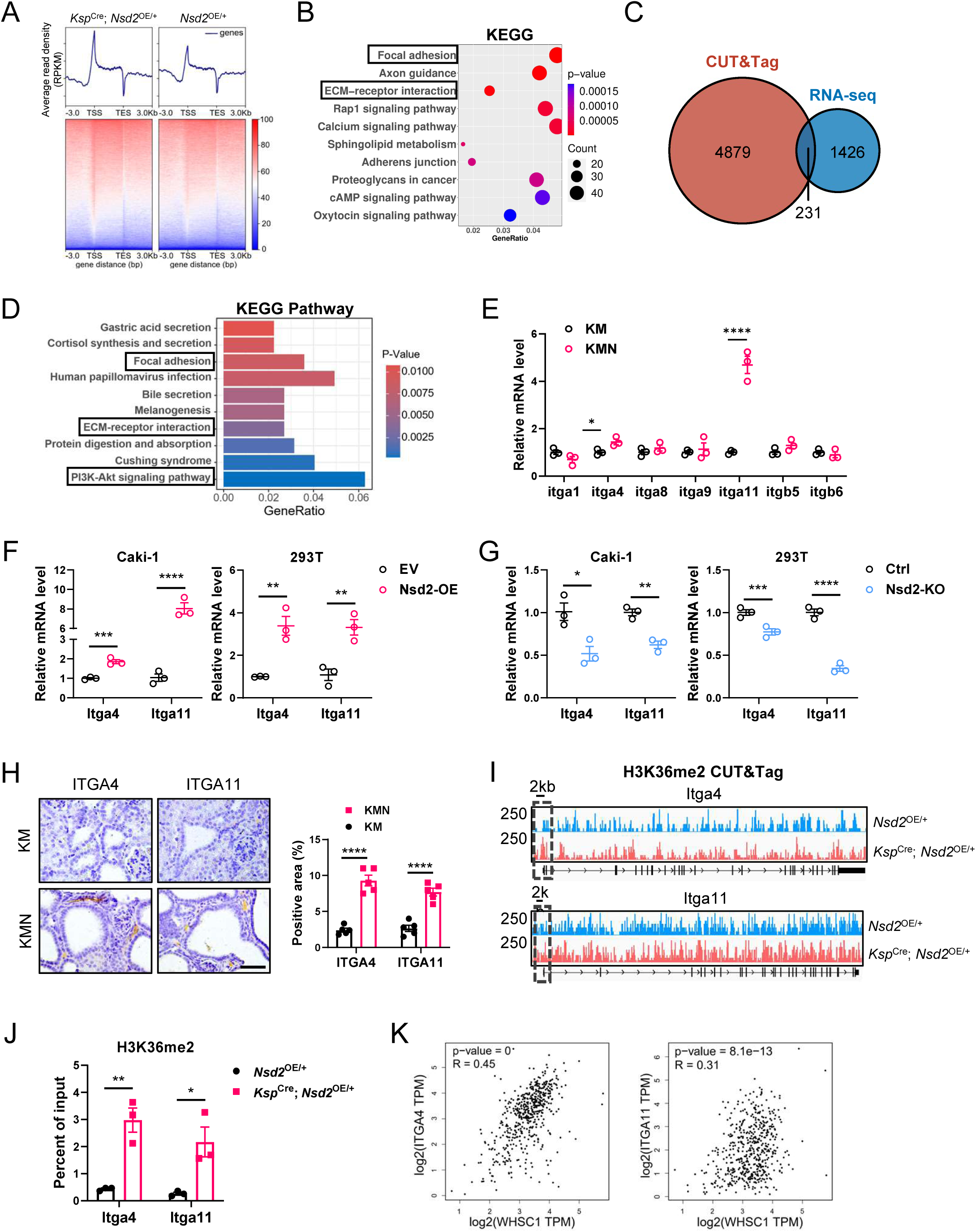
NSD2-mediated H3K36me2 facilitates the expression of integrins. **(A)** Normalized read density of H3K36me2 CUT&Tag signals of *Ksp*^Cre^; *Nsd2*^OE/+^ and *Nsd2*^OE/+^ mice at 12 weeks from 3 kb upstream of the TSS to 3 kb downstream of the TES. **(B)** KEGG pathway analysis of gene expression changes in CUT&Tag data. **(C)** Venn diagram showing the number of genes harboring H3K36me2 binding and the differential expression genes determined by RNA-seq data. **(D)** The KEGG pathway analysis of the overlapping genes in (C). **(E)** RT-qPCR analysis of integrin family mRNA levels of KM and KMN mice (n = 3 per group). Experiments were repeated at least three times, with similar results. **(F)** RT-qPCR analysis of *ITGA4* and *ITGA11* mRNA levels of EV and Nsd2-OE cells (Caki-1 and 293T cells derived, n = 3 per group). Experiments were repeated at least three times, with similar results. **(G)** RT-qPCR analysis of *ITGA4* and *ITGA11* mRNA levels of Ctrl and Nsd2-KO cells (Caki-1 and 293T cells derived, n = 3 per group). Experiments were repeated at least three times, with similar results. **(H)** IHC staining and quantification of ITGA4 and ITGA11 expressions in KM and KMN mice at 12 weeks (n = 5 per group). Experiments were repeated at least three times, with similar results, and representative images are shown. Scale bars: 50 μm. **(I)** Snapshot of H3K36me2 CUT&Tag signals at the *Itga4* and *Itga11* gene in *Nsd2*^OE/+^ and *Ksp*^Cre^; *Nsd2*^OE/+^ mice. **(J)** ChIP-qPCR of *Itga4* and *Itga11* in *Nsd2*^OE/+^ and *Ksp*^Cre^; *Nsd2*^OE/+^ mice. The location of the ChIP primer pairs used is at the promoter area. Experiments were repeated at least three times, with similar results. **(K)** The Pearson product-moment pair-wise gene correlation analysis between *NSD2* and *ITGA4*, *ITGA11* with TCGA database.

Integrins link cells to the ECM, activating FAK to regulate cell adhesion, migration, and signaling transduction, crucial for cellular interaction with the environment (29). Notably, integrin α4 (*Itga4*) and integrin α11 (*Itga11*) were listed among the 231 genes described above, and their upregulations were validated in KMN mice (Fig. 4E). To verify whether NSD2 can regulate the expressions of *ITGA4* and *ITGA11* in vitro, the mRNA levels of *ITGA4* and *ITGA11* were measured in Caki-1 and 293T cells transfected with empty vector and NSD2 overexpression plasmids, respectively. The results showed that *ITGA4* and *ITGA11* were significantly increased in Nsd2-OE cells (Fig. 4F). In addition, *NSD2* was depleted with CRISPR/Cas9 system (referred to as Nsd2-KO), and downregulated *ITGA4* and *ITGA11* expressions were further validated in Nsd2-KO cells (Fig. 4G). According to IHC staining, the protein expression levels of ITGA4 and ITGA11 were also increased in the KMN mice (Fig. 4H).

Direct H3K36me2 occupancies within *Itga4* and *Itga11* gene loci were seen in Genome Browser tracks (Fig. 4I). As expected, we observed a significant increase in H3K36me2 signals in the *Itga4* and *Itga11* gene locus, especially within the promoter regions in NSD2-overexpressing cells (Fig. 4I). We further confirmed the existence of H3K36me2 marks at the promoter of the *Itga4* and *Itga11* genes by ChIP-qPCR, and the signals increased along with NSD2 overexpression (Fig. 4J).

Consistent with these results, there were significant positive correlations between mRNA levels of *NSD2* and that of *ITGA4* and *ITGA11* respectively based on the TCGA database (Fig. 4K). Together, these results indicate that NSD2-mediated H3K36me2 promotes the expression of integrins.

### NSD2 enhances the cell proliferation on the stiff matrix

Considering that FAK signaling plays an important role in regulating cellular behavior resulting from integrin interactions with the ECM (30), PEGDA hydrogel was utilized to mimic the ECM of living tissues.

To elucidate the effect of matrix stiffness on cell behaviors, EV and Nsd2-OE cells were cultured on monomeric collagen type IV-coated PEGDA gels with different stiffness (4 Pa, 0.1 kPa, 0.7 kPa and 20 kPa) (Fig. 5A) (31). One day after plating, both EV and Nsd2-OE cells did not attach to the soft matrix (4 Pa). On modest and stiff matrix (0.1 kPa, 0.7 kPa and 20 kPa), NSD2 induced fibroblast-like morphology and increased cell area, compared with EV cells (Fig. 5B-C). These results indicated that with the increase of matrix stiffness, Nsd2-OE cells displayed much more cell spreading (Fig. 5B-C).

**Fig. 5.**
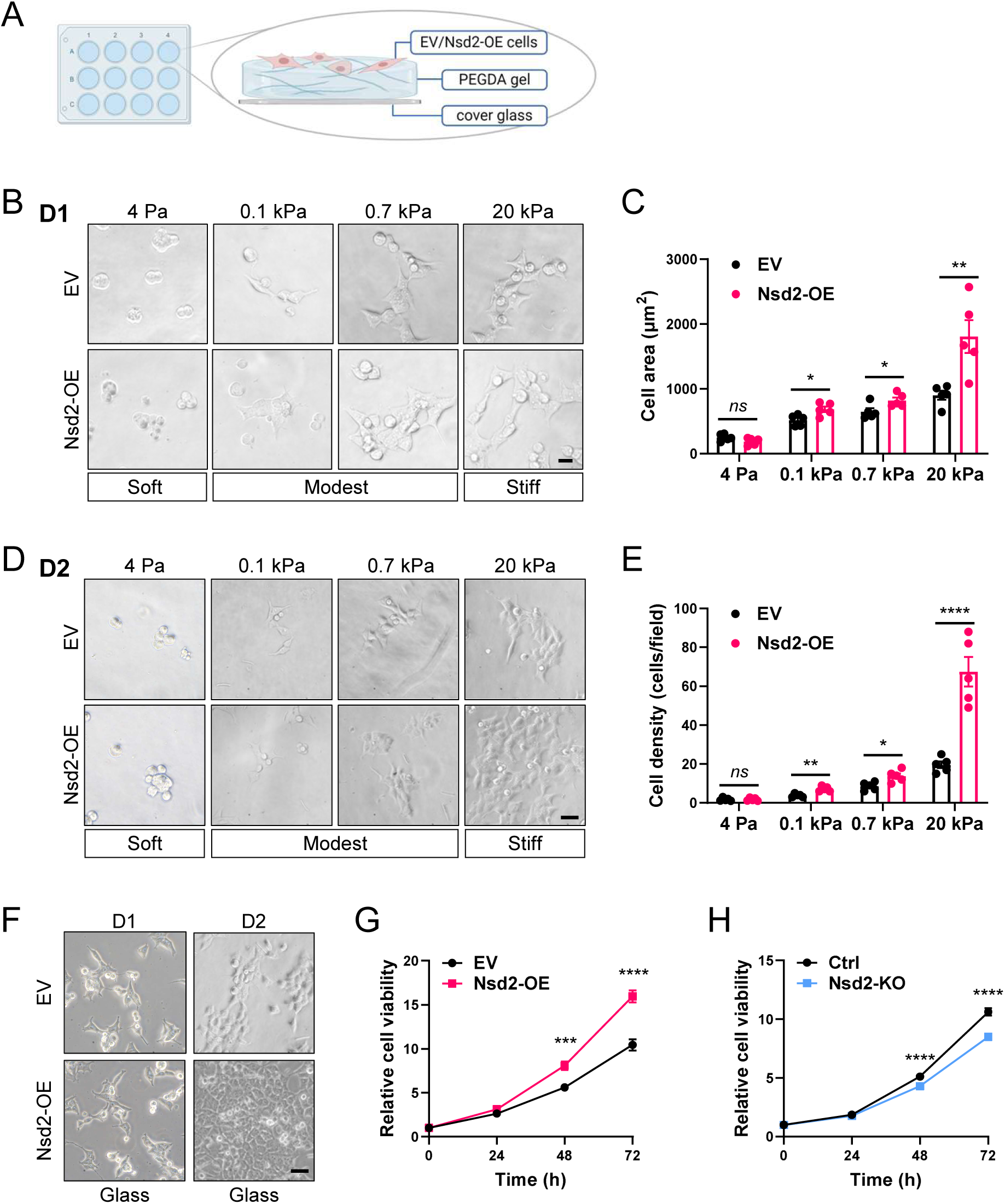
NSD2 enhances the cell proliferation on the stiff matrix. **(A)** Scheme of PEGDA gel for EV and Nsd2-OE cells culture. **(B)** Representative images of EV and Nsd2-OE cells morphology 1 day after plating at 4 Pa, 0.1 kPa, 0.7 kPa, and 20 kPa PEGDA gel. Scale bar: 20 μm. **(C)** Quantification of cell area of EV and Nsd2-OE cells on PEGDA gel 1 day after plating. **(D)** Representative images of EV and Nsd2-OE cells morphology 2 days after plating at 4 Pa, 0.1 kPa, 0.7 kPa, and 20 kPa PEGDA gel. Scale bar: 50 μm. **(E)** Quantification of cell density of EV and Nsd2-OE cells on PEGDA gel 2 days after plating. **(F)** Representative images of EV and Nsd2-OE cells morphology 1 day and 2 days after plating at glass. Scale bar: 50 μm. **(G)** CCK8 assay of EV and Nsd2-OE cells (Caki-1 cells derived). Experiments were repeated at least three times, with similar results. **(H)** CCK8 assay of Ctrl and Nsd2-KO cells (Caki-1 cells derived). Experiments were repeated at least three times, with similar results.

We further analyzed the effect of stiffness on the cell proliferation. Two days after plating, NSD2 significantly promoted the cell proliferation on both modest and stiff matrix, and Nsd2-OE cells displayed much more cell proliferation with the increase of matrix stiffness (Fig. 5D-E). In addition, NSD2 also promoted the cell spreading and proliferation on glass substrate control (Fig. 5F). CCK8 assays confirmed the enhanced cell proliferation of Nsd2-OE cells, and the proliferation of Nsd2-KO cells was significantly decreased compared with the control cells (Fig. 5G-H). Together, NSD2 enhances the cell spreading and proliferation on the stiff matrix.

### Inhibition of the integrin/FAK/AKT signaling alleviates the tumorigenesis of TCRCC

To verify the effect of integrin/FAK signaling in NSD2-mediated tumorigenesis, an α4β1/α4β7 integrin antagonist, Firategrast was utilized in 8-week-old KMN mice (32). The compound was administered by gavage once a day for 21 days under 30 mg/kg dosage (Fig. 6A). As a result, the Firategrast-treated mice exhibited smaller kidney size compared to vehicle-treated mice (Fig. 6B). In addition, smaller cysts areas were observed in Firategrast-treated mice than vehicle-treated mice morphologically (Fig. 6C). IF staining was performed and there are less Col1a1 accumulation around the cysts (Fig. 6D). In further IHC analysis, evidence showed that Firategrast-treated mice exhibited attenuated staining of p-FAK, ITGA4, and ITGA11 (Fig. 6D). The expressions of CK19, AMACR, Vimentin and Ki67 also declined in Firategrast-treated mice (Fig. 6E). These results revealed that inhibiting the integrin signaling reduced the tumorigenesis of TCRCC. To further demonstrate the involvement of ITGA4 in NSD2-overexpression RCC, we depleted *ITGA4* with CRISPR/Cas9 system in Caki-1 cells (referred to as sg-Itga4). Both EV and Nsd2-OE cells were transfected with sg-NC and sg-Itga4 plasmid, respectively. The results indicated that the loss of *ITGA4* declined the proliferation of Nsd2-OE cells but did not affect EV cells. What’s more, knockout of Itga11 (referred to as sg-Itga11) can also decline the proliferation of Nsd2-OE cells (Fig. 6F). Here we conclude that *ITGA4* or *ITGA11* deficiency can attenuate the enhanced proliferation of Nsd2-OE cells (Fig. 6F).

**Fig. 6.**
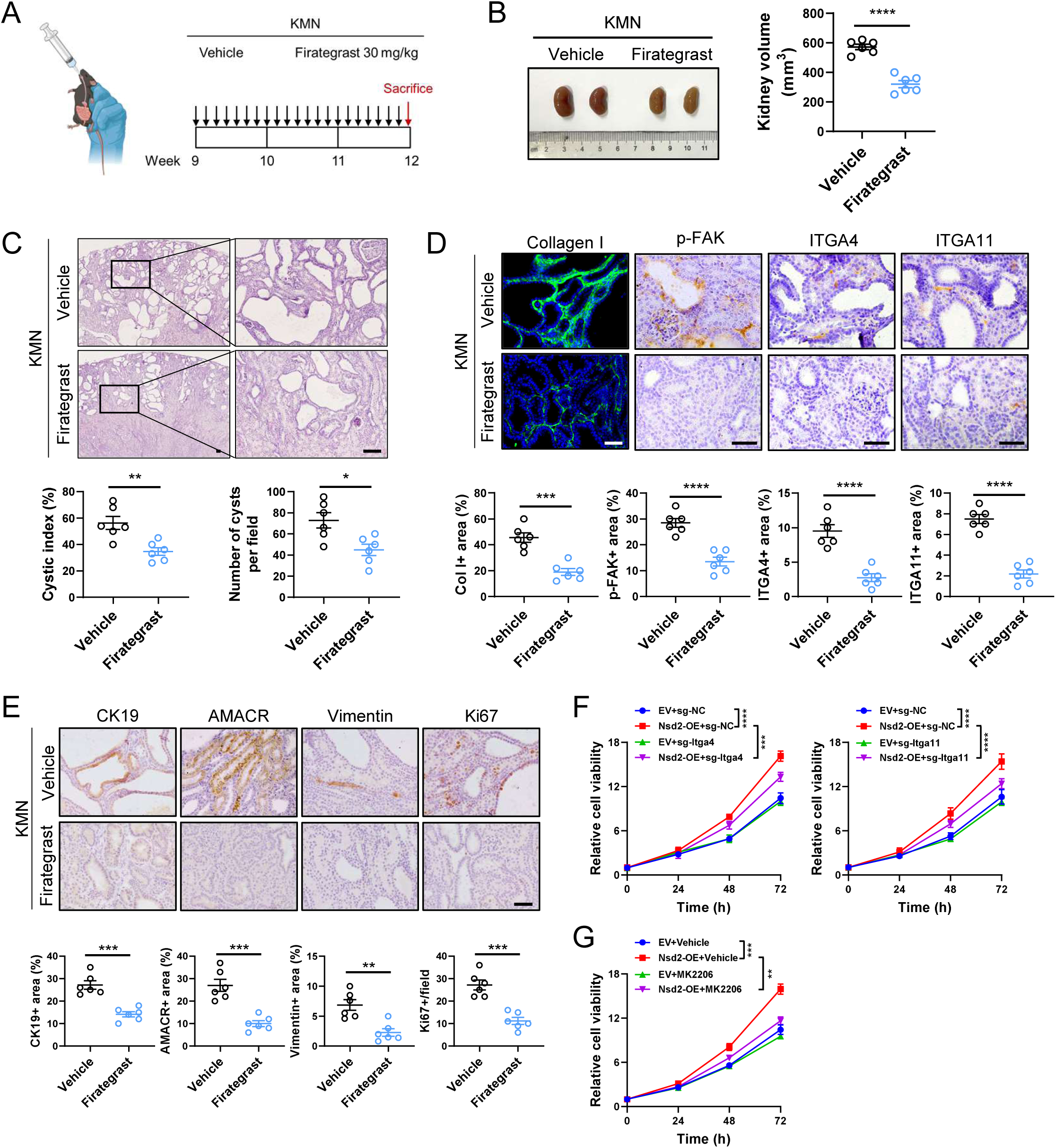
Inhibition of the integrin/FAK/AKT signaling alleviates the tumorigenesis of TCRCC. **(A)** Scheme of treatment given for each gavage. **(B)** Kidney images of KMN mice treated with vehicle or Firategrast for 21 days. Quantification of kidney volume (mm^3^) is on the right (n = 6 per group). **(C)** Representative hematoxylin and eosin images of kidney tissue from KMN mice treated with vehicle or Firategrast for 21 days. Scale bars: 100 μm. Quantification of cystic index (%) and number of cysts per field is shown at the bottom (n = 6 per group). **(D)** IF analysis of Collagen I expression and IHC staining of p-FAK, ITGA4, and ITGA11 in KMN mice treated with vehicle or Firategrast for 21 days. Scale bars: 50 μm. Quantification of Collagen I, p-FAK, ITGA4 and ITGA11 positive area is shown at the bottom (n = 6 per group). **(E)** IHC staining and quantification of CK19, AMACR, Vimentin positive area, and Ki67 positive cells per field from indicated mice (n = 6 per group). Scale bars: 50 μm. **(F)** CCK8 assay of EV and Nsd2-OE cells transfected with the indicated plasmid, respectively. Experiments were repeated at least three times, with similar results. **(G)** CCK8 assay of EV and Nsd2-OE cells treated with MK2206 or vehicle, respectively. Experiments were repeated at least three times, with similar results.

**Fig. 7.**
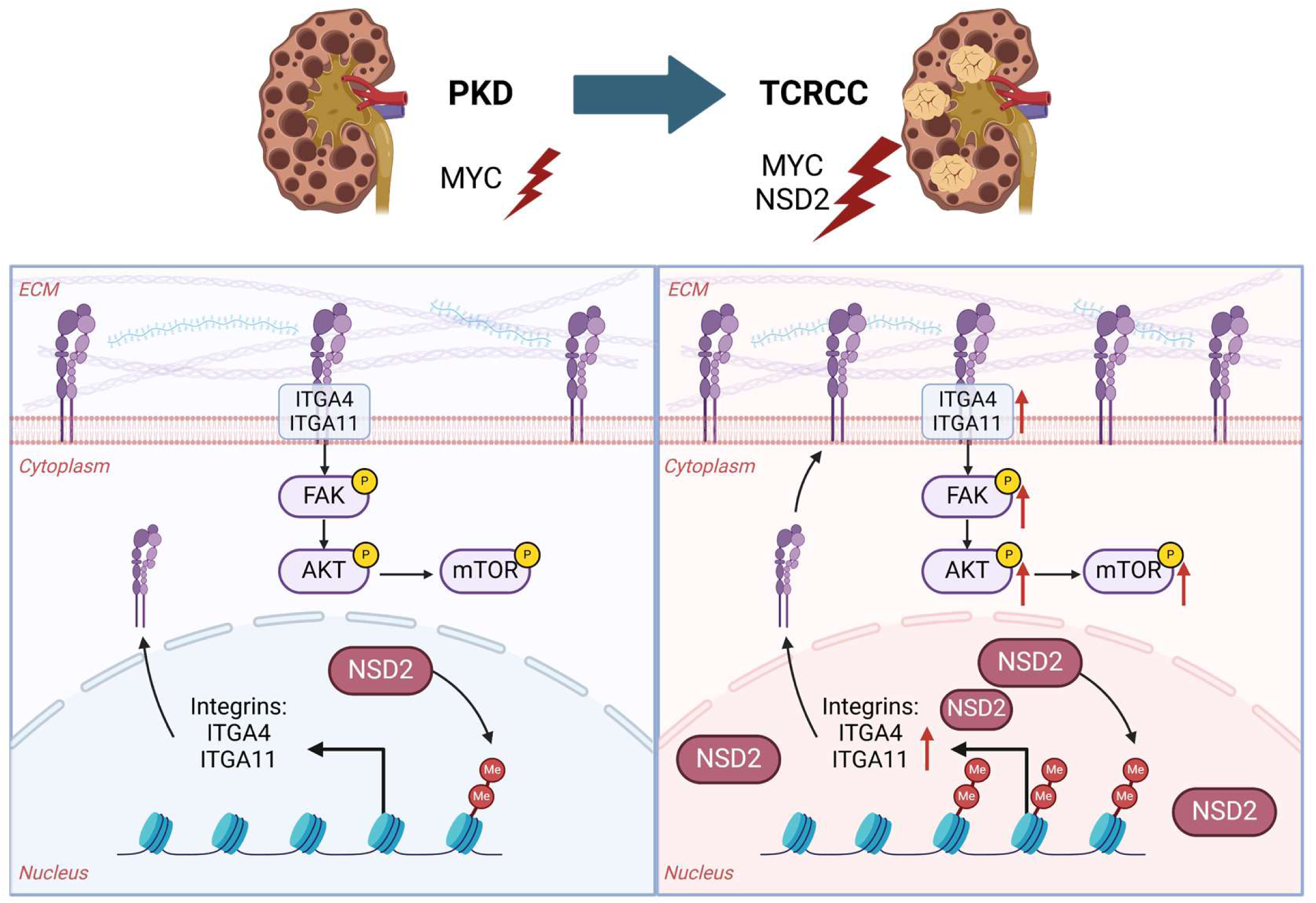
NSD2 promotes the transition from PKD to TCRCC through activating the integrin/FAK/AKT signaling.

As we mentioned above, FAK can lead to TCRCC tumorigenesis through activating AKT signaling pathway. Therefore, MK-2206, the inhibitor of AKT pathway(33), was utilized to verify the function of AKT in Nsd2-OE cells. The results of CCK8 assay showed that with the treatment of MK-2206, the proliferation of Nsd2-OE cells was significantly suppressed. (Fig. 6G). Collectively, these results demonstrated that the inhibition of integrin/FAK/AKT signaling alleviates the tumorigenesis of TCRCC.

## Discussion

TCRCC is an extremely rare subtype of renal cell carcinoma that was officially included as an independent subtype of renal cell carcinoma by the World Health Organization (WHO) in 2016 (34). Less than 100 cases of pure TCRCC have been reported in the literature (34). As a rare type of kidney tumor, TCRCC has relatively limited clinical and pathologic information in the literature, and no animal model is available. In this study, we generated a mouse model with kidney-specific overexpression of MYC and NSD2 (KMN). We found that the KMN mice developed a significant cystic phenotype at only 6 weeks of age, and gradually progressed to TCRCC at 12 weeks of age (Fig. 4D-E). Therefore, the *Myc* and *Nsd2* double-transgenic mice that we established may serve as a novel autochthonous genetically engineered mouse model of TCRCC for preclinical research.

Our previous study has revealed that loss of SETD2 promotes the transition from PKD to ccRCC by activating the Wnt/β-catenin pathway (35), which results in the disordered proliferation of renal tubular epithelial cells. In current study, NSD2 promotes the transition from *Myc*-induced PKD to TCRCC. Our results demonstrated that NSD2-mediated TCRCC in mice is featured by closely packed well-formed tubules and cysts. These cysts are surrounded by abundant ECM (Fig. 3E), and no papilla or solid sheets of clear cells are seen within the tubules or cysts. Notably, NSD2 enhances the cell spreading and proliferation on the stiff matrix in vitro (Fig. 5). These results collectively suggest a phenomenon of durotaxis of NSD2-overexpressing cells. Previous studies have shown that FAK is crucial in durotaxis, which is a cellular phenomenon where cells migrate along gradients of ECM stiffness, typically towards areas of greater stiffness (36). Changes in ECM stiffness are sensed by these focal adhesions, leading to the activation of FAK and subsequent signaling pathways that govern cell migration and cytoskeletal organization (37,38). Moreover, a previous study found that FAK knockdown has been shown to inhibit the directionality that fibroblasts normally display when undergoing durotaxis on a stiffness gradient (39). The durotaxis due to FAK signaling activation might lead to the TCRCC phenotype of NSD2-overexpressing mice, and the underlying mechanism needs to be further studied.

In summary, our results demonstrate that NSD2 promotes the transition from *Myc*-induced PKD to TCRCC through integrin/FAK/AKT pathway. In addition, NSD2 enhances the cell proliferation on the stiff matrix. Inhibition of the integrin/FAK signaling alleviates the tumorigenesis of TCRCC. Our study advances the understanding of the pathogenesis of TCRCC and opens therapeutic opportunities for TCRCC patients with NSD2 amplification or high expression by treatment with integrin/FAK/AKT inhibitors.

## Acknowledgments

This work was supported by funds from National Key R&D Program of China (2022YFA1302704 to L.L. and W.-Q.G.), National Natural Science Foundation of China (82372604, 82073104 to L.L., U23A20441 to W.-Q.G.), Science and Technology Commission of Shanghai Municipality (21JC1404100 to W.-Q.G.) and 111 project (no. B21024). We thank Genefund Biotech (Shanghai, China) for assistance in the data analysis.

## Author contributions

L.L. designed experiment and interpreted data; W.F and N.L. performed most of the experiments; C.L, H.R, Z.C., W.Z., Y.X., R.A. and Z.W. assisted in some experiments; W.-Q.G. assisted in some discussion; W.F., N.L. and L.L. wrote the manuscript; L.L. provided the overall guidance.

## Data sharing statement

The sequencing data sets have been deposited in the National Center for Biotechnology Information (NCBI)’s Gene Expression Omnibus (GEO). RNA-Seq raw data have been deposited in GEO under accession number: GSE266614. CUT-Tag raw data have been deposited in GEO under accession number: GSE264634.

