## Supplementary figures for "NSD2 promotes cell durotaxis and drives the transition from PKD to tubulocystic renal cell carcinoma through integrin/FAK/AKT signaling"

### Figure S1

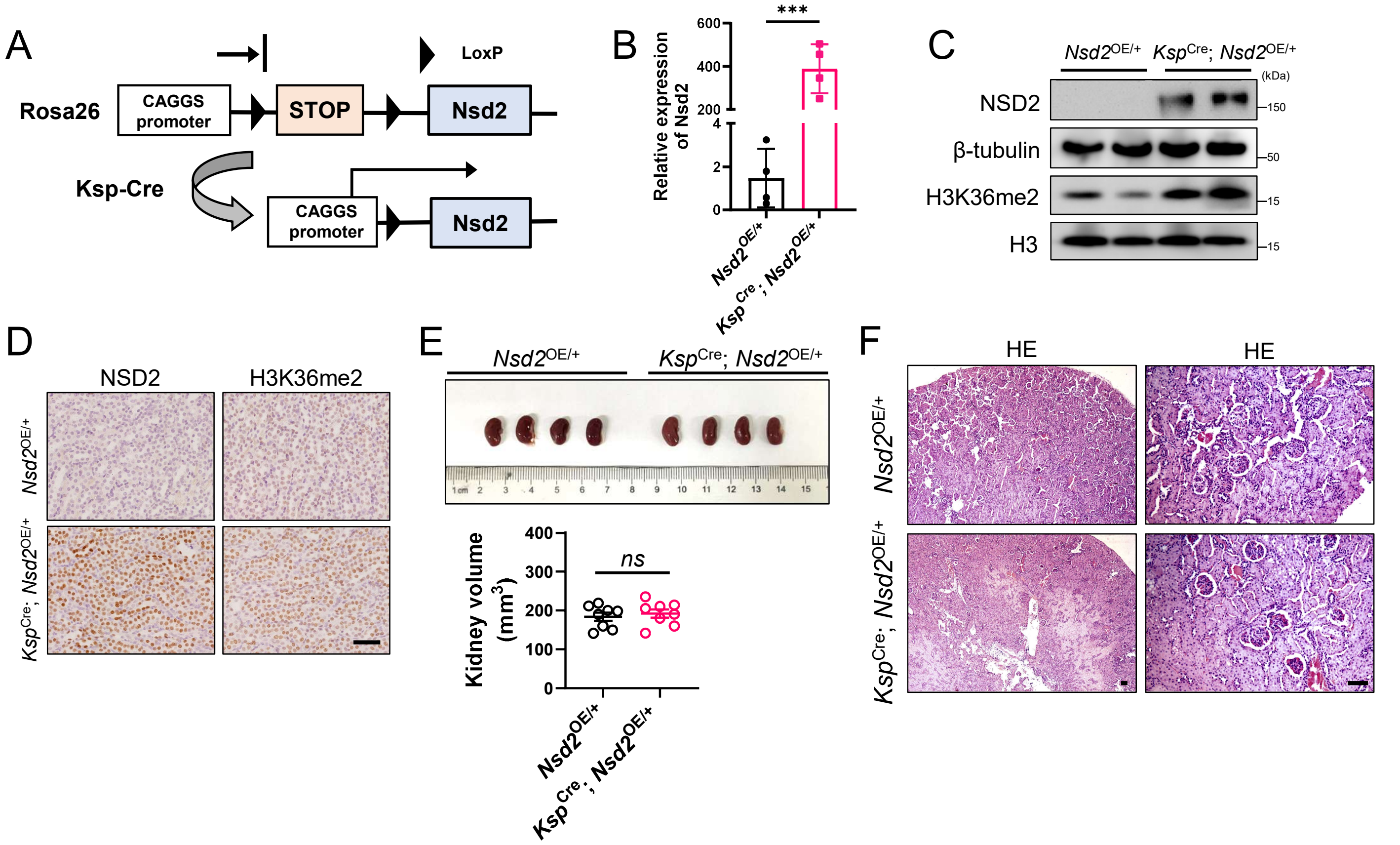


**Fig. S1. Histological examination of *Nsd2*$^{\mathbf{OE/+}}$ and *Ksp*$^{\mathbf{Cre}}$; *Nsd2*$^{\mathbf{OE/+}}$ mice at 12 months of age.**

(A) Construction and breeding strategy of *Ksp*$^{Cre}$; *Nsd2*$^{OE/+}$ mice.

(B) RT-qPCR analysis of *Nsd2* mRNA level in kidney tissues of *Nsd2*$^{OE/+}$ and *Ksp*$^{Cre}$; *Nsd2*$^{OE/+}$ mice. Experiments were repeated at least three times, with similar results.

(C) Western blotting analysis of NSD2 and H3K36me2 levels in kidney tissues of *Nsd2*$^{OE/+}$ and *Ksp*$^{Cre}$; *Nsd2*$^{OE/+}$ mice. Experiments were repeated at least three times, with similar results.

(D) IHC analysis of NSD2 and H3K36me2 levels in kidney tissues of *Nsd2*$^{OE/+}$ and *Ksp*$^{Cre}$; *Nsd2*$^{OE/+}$ mice. Scale bars: 50 μm.

(E) Kidney images of *Nsd2*$^{OE/+}$ and *Ksp*$^{Cre}$; *Nsd2*$^{OE/+}$ mice in 12 month. Quantification of kidney volume ($mm^3$) is at the bottom.

(F) Representative hematoxylin and eosin images of kidney tissue from 12-month *Nsd2*$^{OE/+}$ and *Ksp*$^{Cre}$; *Nsd2*$^{OE/+}$ mice. Scale bars: 50 μm.

### Figure S2

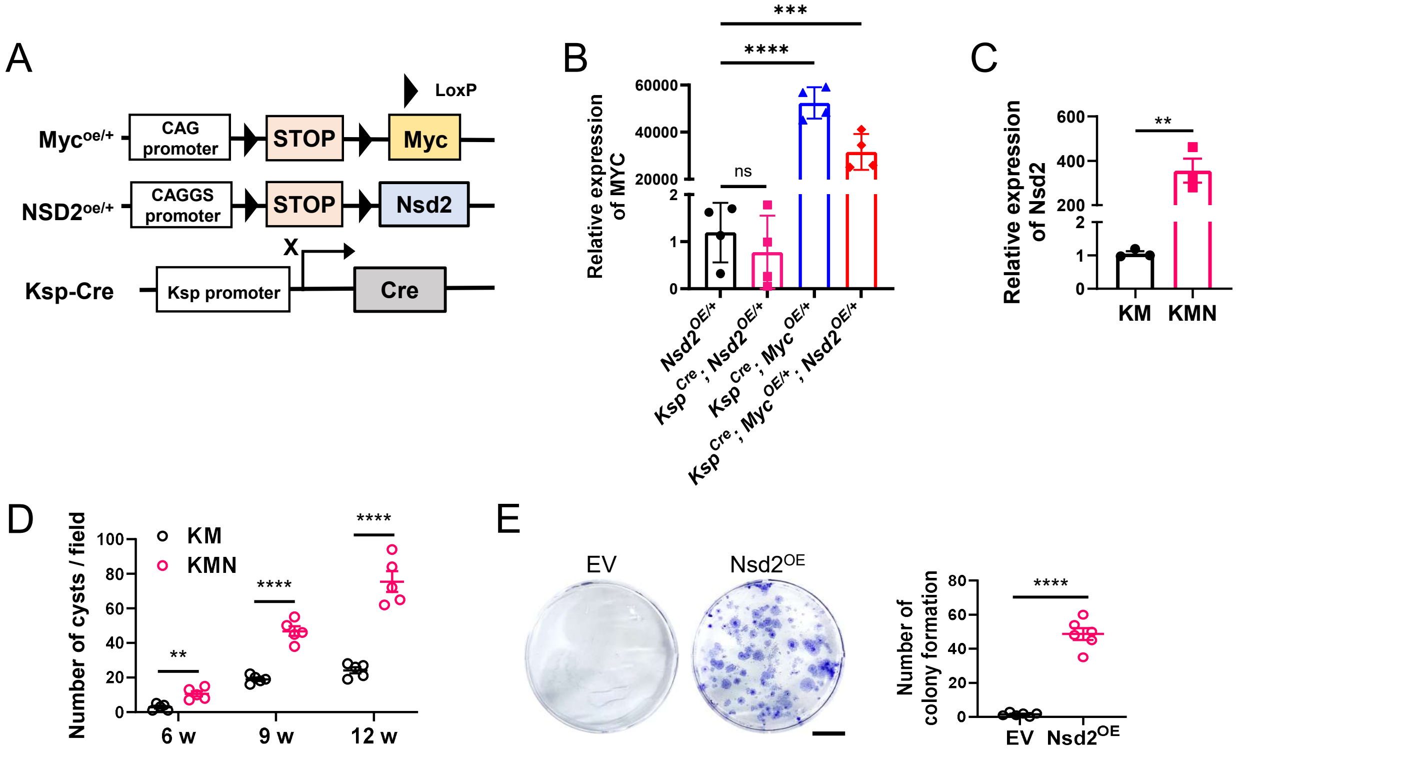


**Fig. S2. Construction of *Ksp*$^{Cre}$; *Myc*$^{OE/+}$; *Nsd2*$^{OE/+}$ mice.**

(A) Construction and breeding strategy of *Ksp*$^{Cre}$; *Myc*$^{OE/+}$; *Nsd2*$^{OE/+}$ mice (KMN).

(B) RT-qPCR analysis of *Myc* mRNA level in kidney tissues of *Nsd2*$^{OE/+}$, *Ksp*$^{Cre}$; *Nsd2*$^{OE/+}$, *Ksp*$^{Cre}$; *Myc*$^{OE/+}$ and *Ksp*$^{Cre}$; *Myc*$^{OE/+}$; *Nsd2*$^{OE/+}$ mice. Experiments were repeated at least three times, with similar results.

(C) RT-qPCR analysis of *Nsd2* mRNA level in kidney tissues of KM and KMN mice. Experiments were repeated at least three times, with similar results.

(D) Quantification of cysts number per field in kidney tissue from indicated mice at 6, 9, and 12 weeks (n = 5 per group).

(E) Representative images and quantification of colony formation in indicated cells (n = 6 per group). Scale bars: 5 mm.

### Figure S3

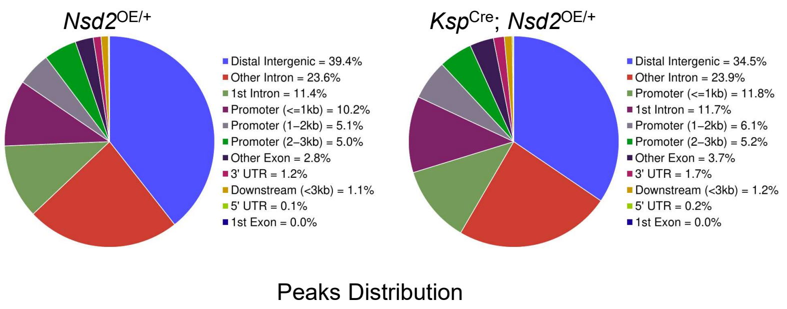


**Fig. S3. Analysis of the occupancy of H3K36me2 CUT-Tag peaks.**

Analysis of the occupancy of H3K36me2 CUT-Tag peaks of *Nsd2*$^{OE/+}$ and *Ksp*$^{Cre}$; *Nsd2*$^{OE/+}$ mice. Different genomic annotations are marked at indicated colors.
